# Contribution of sensory encoding to measured bias

**DOI:** 10.1101/444430

**Authors:** Miaomiao Jin, Lindsey L. Glickfeld

**Author notes:** Corresponding Author and Lead Contact: Lindsey Glickfeld, Department of Neurobiology, Duke Medical School, 311 Research Drive, BRB-401F, Durham, NC 27710.

## Abstract

Perceptual decision-making is a complex process that involves sensory integration followed by application of a cognitive threshold. Signal detection theory (SDT) provides a mathematical framework for attributing the underlying neurobiological processes to these distinct phases of perceptual decision-making. In particular, SDT reveals the sensitivity (d’) of the neuronal response distributions and the bias (c) of the decision criterion, which are commonly thought to reflect sensory and cognitive processes, respectively. However, neuronal representations of bias have been observed in sensory areas, suggesting that some changes in bias are due to effects on sensory encoding. To directly test whether sensory encoding can influence bias, we optogenetically manipulated neuronal excitability in primary visual cortex (V1) during a detection task. Increasing excitability in V1 significantly decreased behavioral bias, while decreasing excitability had the opposite effect. To determine whether this change in bias is consistent with the effects on sensory encoding, we made extracellular recordings from V1 neurons in passively viewing mice. Indeed, we found that optogenetic manipulation of excitability shifted the neuronal bias in the same direction as the behavioral bias, despite using a fixed artificial decision criterion to predict hit and false alarm rates from the neuronal firing rates. To test the generality these effects, we also manipulated the quality of V1 encoding by changing stimulus contrast or inter-stimulus interval. These stimulus manipulations also resulted in consistent changes in bias measured both behaviorally and neuronally. Thus, changes in sensory encoding are sufficient to drive changes in bias measured using SDT.

## Introduction

Perceptual decision-making is a multi-step process though which sensory information about the external world is first transformed into a neuronal code and then used to make a behavioral choice. In this process, both sensory encoding and the cognitive aspects of the decision-making process are critical factors that determine the final choice (Gold and Shadlen, 2007; Carandini and Churchland, 2013; Romo and de Lafuente, 2013; Hanks and Summerfield, 2017).

Efforts to dissect the relative contribution sensory and cognitive processes to decision-making often take advantage of signal detection theory (SDT), a classical and widely used method that allows inference of the underlying neuronal response distributions and decision criteria from behavioral measures (Green and Swets, 1966). In particular, SDT allows the use of hit and false alarm (FA) rates to extract two aspects of the perceptual decision: sensitivity (d’) and bias (c). Measures of sensitivity allow inference of the separability of the underlying neuronal activity evoked in response to targets and distractors. Thus, this measure is thought to reflect the quality of encoding in sensory circuits that provide input to the decision-making circuits (Bashinski and Bacharach, 1980; Bennett et al., 2013; Pinto et al., 2013; Luo and Maunsell, 2015; Jurjut et al., 2016; Ni et al., 2017). On the other hand, bias measures the overall tendency to classify the stimulus as a target or distractor. Thus, it can reflect the subject’s decision criterion. In fact, c is often used synonymously with “criterion” and is therefore commonly thought to reveal cognitive contributions to the decision-making process and involve areas downstream of sensory cortex (McDonald et al., 2000; Grove et al., 2012; Jones et al., 2015; Crapse et al., 2017; de Gee et al., 2017; Luo and Maunsell, 2018; van Vugt et al., 2018).

However, neuronal correlates of bias have also been identified in sensory cortical areas. Human neuroimaging experiments have found a strong correlation between the strength of representation of prior information (such as expected stimulus features or locations) in sensory areas and the strength of behavioral bias (White et al., 2012; Kok et al., 2013; Vintch and Gardner, 2014). Similarly, spontaneous fluctuations in the excitability of sensory cortical areas correlate with spontaneous fluctuations in behavioral bias (Iemi et al., 2017). These data suggest that activity in sensory areas can influence behavioral bias. However, it is not clear whether this is due to a direct effect of sensory encoding on bias or the result of a bidirectional interaction between sensory and cognitive systems.

In fact, there is a clear mathematical explanation for how changes in sensory encoding can alter behavioral bias (Witt et al., 2015). Since bias is always measured relative to the optimal criterion (**Figure 1a**), changes in sensory encoding that shift the optimal criterion have the potential to result in changes in measured bias. This happens any time that changes in the responses to the target and distractor are not opposite and proportional. Thus, many manipulations that alter sensory encoding, ranging from adaptation to attention, might be expected to cause changes in bias in addition to sensitivity, even in the absence of a cognitive contribution.

**Figure 1.**
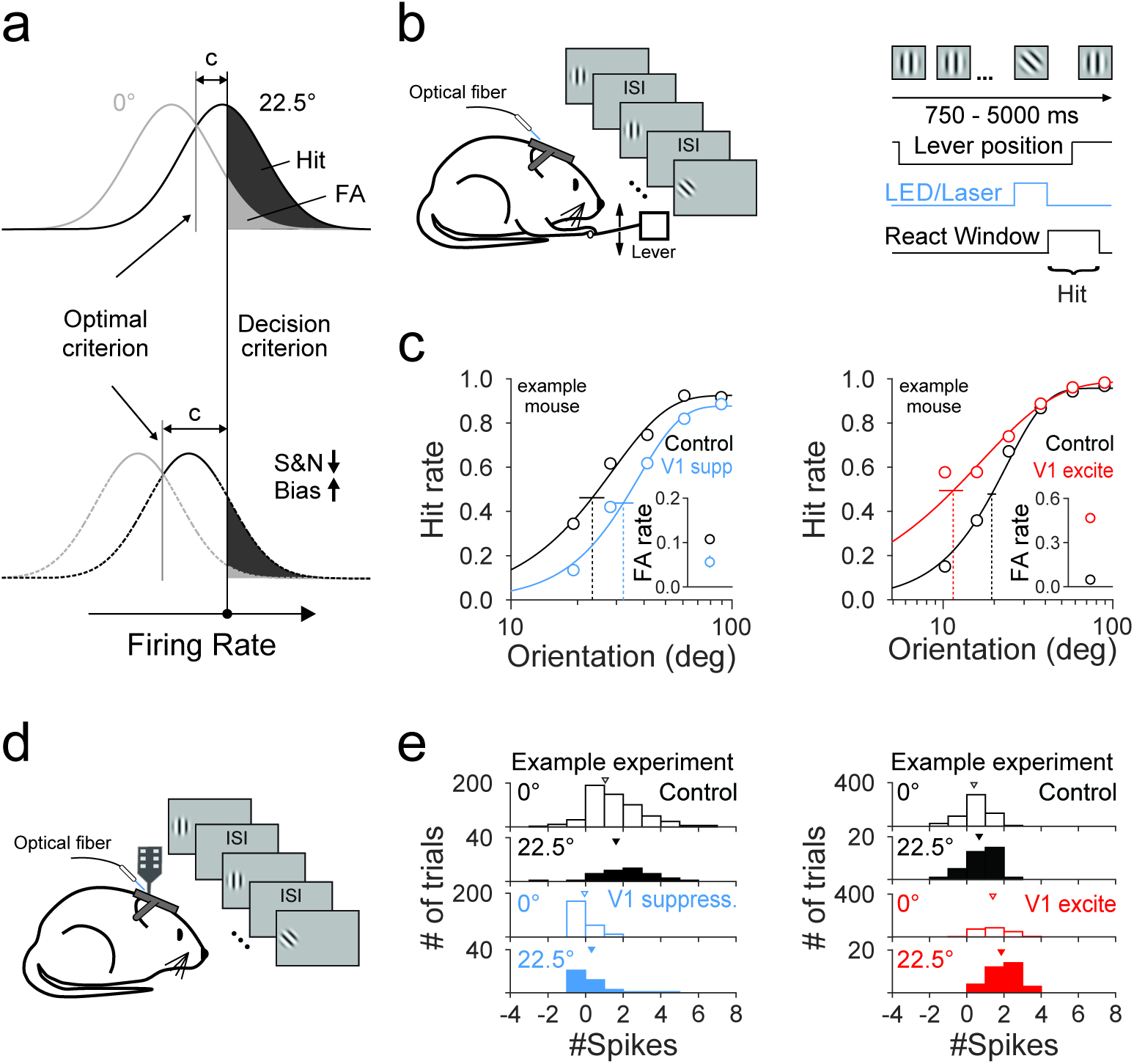
Optogenetically suppressing or exciting V1 decreases or increases both signal and noise distributions in an orientation discrimination task. (**a**) Schematic of effect of shifting signal and noise distributions on bias measured using signal detection theory. Top: distributions of target (22.5°, solid black) and distractor (0°, solid gray) responses. Note that bias (c) is measured as the distance between the actual (black vertical line) and optimal (c=0, gray vertical line) criterion. Bottom: manipulations that decrease both the target and distractor distributions shift the optimal criterion to the left, and therefore result in an increase in bias. (**b**) Schematic of behavior setup and trial progression. Blue light is turned on for a single target or distractor presentation on each trial. V1 suppression (blue) and excitation (red) is achieved via optogenetically driving PV+ or VGAT+ neurons and Emx1+ neurons respectively. (**c**) Hit rate and FA rate (inset) for control (black) and V1 suppression (blue, left) or excitation (red, right) for one example mouse each. Hit rates are fit with a Weibull function; vertical dotted lines are threshold, error is 95% confidence interval. (**d**) Schematic of extracellular recording setup. Stimuli are presented as in **b**. (**e**) Distributions of spikes summed across a simultaneously recorded population in response to distractor (0°, open bars) and target (22.5°, filled bars) stimuli on control trials (black) and during V1 suppression (blue, n=17 cells, left) or excitation (red, n=16 cells, right) for one example experiment each. Triangles show the mean of the distribution.

To directly test whether changes in sensory encoding are sufficient to affect bias, we trained mice on an orientation discrimination task in which we could 1) measure hit and FA rate to calculate bias and sensitivity and 2) control the neuronal responses to both targets and distractors. Altering responses to targets and distractors through either direct optogenetic manipulation of neurons in primary visual cortex (V1) or manipulation of visual stimulus properties results in a reliable change in behavioral bias with relatively little impact on sensitivity. Further, electrophysiological recordings from neurons in V1 during each of these manipulations also revealed a strong effect on bias in the same direction as during behavior. Thus, changes in bias can be driven by changes in either cognitive factors or sensory encoding, and the lack of a change in sensitivity does not preclude a change in sensory encoding.

## Results

To explore whether purely sensory changes can affect measured bias in perceptual decision-making, we designed an orientation discrimination task to allow measures of hit and false alarm (FA) rate (**Figure 1b**). In this task, a head-fixed mouse presses a lever to initiate trials and releases it to report a target orientation. Each trial begins with the repeated presentation of at least two (and up to nine) iso-oriented gratings (‘distractors’, 100 ms duration) followed by a counterclockwise change in orientation relative to the distractor (‘target’, range: 9-90°; **Figure 1c**). If the mouse releases the lever within a window 200-550 ms following the onset of the target stimulus, it is considered a hit; if the mouse releases the lever within the same window following a distractor stimulus, it is considered a FA. Thus, we can use these behavioral measures to calculate sensitivity and bias using SDT (Green and Swets, 1966).

In addition to being appropriate for making measurements of SDT, this task has a couple of additional advantages. First, the mice can perform the task at a high level of proficiency with low lapse rates (0.053±0.008; range 0.003-0.107; n=14 mice), FA rates (0.048±0.004; range 0.032-0.098; n=14 mice) and threshold for orientation discrimination (25.2°±1.3°; range 14.2°-32.0°; n=14 mice). Thus, there are minimal concerns about changes in motivational state or arousal that could influence our measures of bias. Second, we have a good idea of how neuronal activity in primary visual cortex (V1) is used to perform the behavior (Jin et al., 2018). Namely, the decision-making circuits sum V1 spike rates, with particular weight on the neurons that prefer targets. Thus, the decision variables and decision criterion are in units of firing rate, and manipulations that coincidently alter firing rates in response to distractors and targets will change the optimal criterion and therefore induce a change in measured bias (**Figure 1a**).

### Direct suppression and activation of V1 alters both behavioral and neuronal measures of bias

To directly test the contribution of sensory encoding in V1 to measures of bias, we optogentically manipulated the firing rates (FR) of V1 neurons. We virally or genetically expressed excitatory opsins (ChR2 or Chronos) in either inhibitory or excitatory neurons using transgenic mouse lines (PV::Cre or VGAT-ChR2 and EMX1::Cre). We then used blue light to suppress or excite V1 neurons specifically during presentation of targets or distractors either during performance of the orientation discrimination task (**Figure 1b-c**) or in passively viewing mice (**Figure 1d-e**). Indeed, extracellular recordings from V1 neurons reveal that optogenetic activation of inhibitory neurons significantly reduces neuronal responses to both targets near the animals’ discrimination threshold and distractors (FR changes by V1 suppression: 22.5°: −6.6±1.2 Hz, p<10^-9^; 0°: −5.5±1.2 Hz, p<10^-10^; n=70 cells; Wilcoxon signed rank test; an example experiment in **Figure 1e** and all cells in **Figure S1c**), while activation of excitatory neurons increases visually driven responses (FR changes by V1 excitation: 22.5°: 2.6±0.4 Hz, p<10^-8^; 0°: 2.5±0.3 Hz, p<10^-11^; n=83 cells; Wilcoxon signed rank test; an example experiment in **Figure 1e** and all cells in **Figure S1g**). Moreover, the waveform shapes between control and optogenetic manipulations remain relatively similar (correlation coefficient: control vs. V1 suppression: 0.993 ± 0.001; control vs. V1 excitation: 0.997±0.001; **Figure S1b,f**). Importantly, these effects are largely selective for the targeted stimulus as we see little to no effect on stimuli (Stim_N_) for which the preceding stimulus (Stim_N-1_) was optogenetically manipulated (FR changes by V1 suppression: 22.5°: −1.2±0.6 Hz, p=0.02; 0°: −0.5±0.4 Hz, p=0.50; FR changes by V1 excitation: 22.5°: −0.2±0.3 Hz, p=0.58; 0°: −0.03±0.11 Hz, p=0.79; Wilcoxon signed rank test; **Figure S1d,h**).

Consistent with activation of inhibitory interneurons reducing firing rates, and therefore the decision variable, we find that optogenetic suppression of activity in V1 reduces behavioral hit rate (22.5° target: V1 suppression vs. Control: p<0.005; n=4 mice; paired t-test; **Figure 2a**) and FA rate (V1 suppression vs. Control: p<0.05; n=4 mice; paired t-test). These associated changes in both hit and FA rate often reflect changes in bias (c) measured by SDT. Indeed, using SDT we find a significant increase in measured bias (c for 22.5° target: V1 suppression vs. Control: p<0.005; paired t-test; **Figure 2b**) and a slight decrease in sensitivity (d’ for 22.5° target: V1 suppression vs. Control: p=0.05; paired t-test). Conversely, optogenetic excitation of V1 increases behavioral hit rates (22.5° target: V1 excitation vs. Control: p<0.01; n=4 mice; paired t-test; **Figure 2a**) and FA rate (V1 excitation vs. Control: p<0.01; n=4 mice; paired t-test), resulting in a decrease in measured bias (c for 22.5° target: V1 excitation vs. Control: p<0.005; paired t-test; **Figure 2b**) and a slight decrease in sensitivity (d’ for 22.5° target: V1 excitation vs. Control: p=0.05; paired t-test).

**Figure 2.**
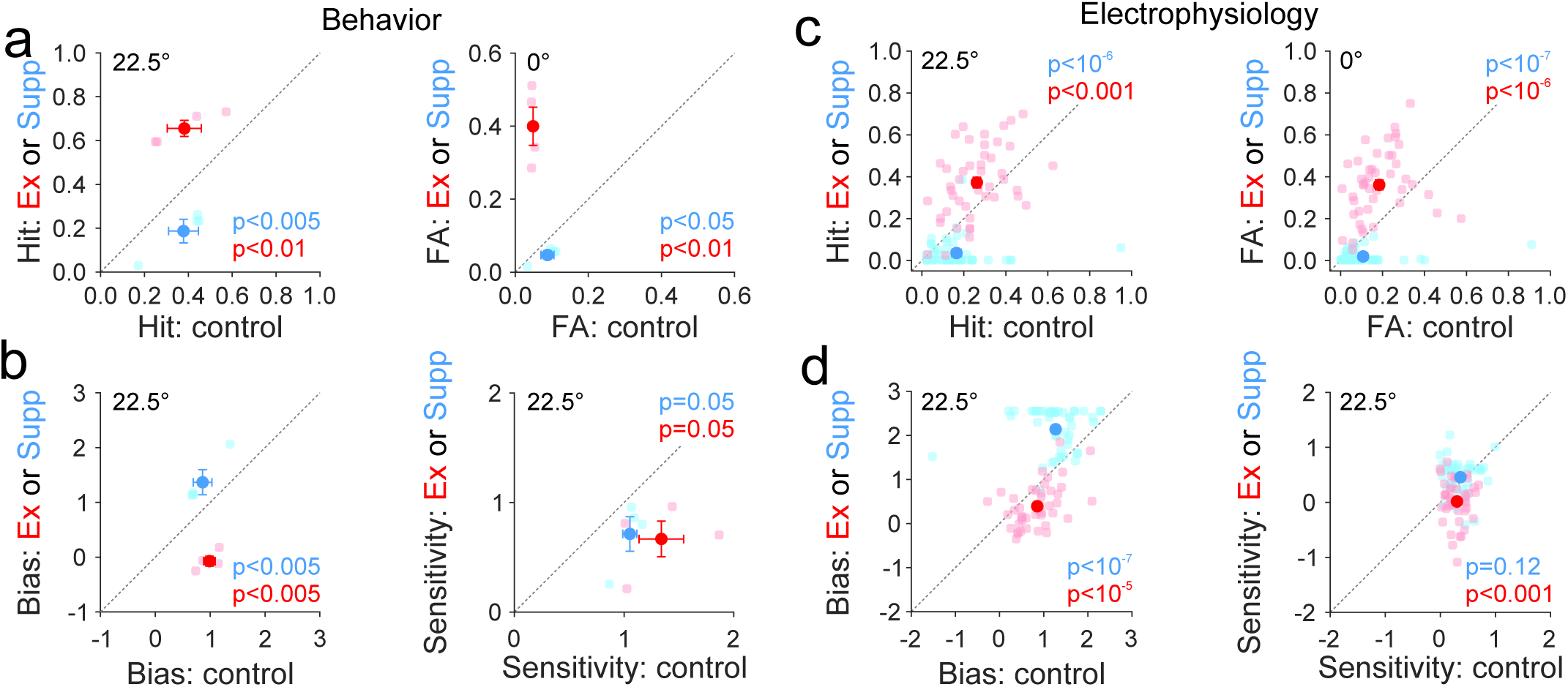
Suppressing or exciting V1 increases or decreases behavioral and neuronal bias. (a) Comparison of the hit (22.5°, left) and FA rate (0°, right) between control and V1 suppression (blue, n=4 mice) or excitation (red, n=4 mice). Light colors are individual mice and dark colors are the mean of the population. Error bars are SEM across mice. **(b)** Same as **a**, for bias (left) and sensitivity (right) at 22.5°. (**c**) Same as **a**, for predicted hit (22.5°, left) and FA rate (0°, right) from neuronal responses using a fixed criterion for each cell (see Methods). (**d**) Predicted bias (left) and sensitivity (right) using the predicted hit and FA rate in **c**. Extreme values of hit and FA rate were corrected (see Methods). Error bars are SEM across cells (V1 suppression-blue: n=47 cells, 3 mice; V1 excitation-red, n=45 cells, 3 mice).

To test whether the changes in firing rate can qualitatively account for the changes in behavioral bias, we used the neuronal data to directly measure bias by applying an artificial decision criterion. We set the criterion to optimally discriminate the optogenetically suppressed distractors (0°) from targets in the control condition (22.5°) in each cell, and then used the distributions of neuronal responses to calculate hit and FA rate across conditions. Suppressing neuronal activity in V1 decreases the predicted hit rate (22.5° target: V1 suppression vs. Control: p<10^-6^; n=47 cells; Wilcoxon signed rank test; **Figure 2c**) and FA rate (V1 suppression vs. Control: p<10^-7^; n=47 cells; Wilcoxon signed rank test) leading to an increase in measured bias (c for 22.5° target: V1 suppression vs. Control: p<10^-7^; Wilcoxon signed rank test; **Figure 2d**) without significantly changing the sensitivity (d’ for 22.5° target: V1 suppression vs. Control: p=0.12; Wilcoxon signed rank test). Conversely, activating neuronal activity in V1 increases predicted hit (22.5° target: V1 excitation vs. Control: p<0.001; n=45 cells; Wilcoxon signed rank test; **Figure 2c**) and FA rate (V1 excitation vs. Control: p<10^-6^; n=45 cells; Wilcoxon signed rank test), resulting in a decrease in measured bias (c for 22.5° target: V1 excitation vs. Control: p<10^-5^; Wilcoxon signed rank test; **Figure 2d**) and sensitivity (d’ for 22.5° target: V1 excitation vs. Control: p<0.001; Wilcoxon signed rank test). Thus, our electrophysiology data shows that manipulating excitability of neurons in V1 is sufficient to alter bias even in the absence of a flexible decision criterion. Moreover, the neuronal and behavioral changes in bias are in the same direction, suggesting that the changes in sensory encoding could be responsible for the changes in behavioral bias.

### Manipulation of stimulus contrast affects measures of bias

Optogenetic tools allow for the direct manipulation of firing rates, however any manipulation that coincidently increases or decreases firing rates in response to targets and distractors are predicted to impact measures of bias. For instance, neurons in V1 have monotonic contrast-response functions (Gao et al., 2010), and therefore decreasing stimulus contrast should decrease firing rates in response to both targets and distractors, shifting the optimal criterion to lower stimulus values. Thus, we modified our orientation discrimination task to vary stimulus contrast (30%, 50% and 70%) on a presentation-by-presentation basis (**Figure 3a**). Extracellular recordings confirm that manipulation of contrast significantly affected firing rates in response to both targets (22.5°: FR changes from 70% to 30%: −4.5±0.7 Hz, p<10^-7^; n=92 cells; Friedman test (p<10^-8^) with post-hoc Tukey HSD test; **Figure S3a**) and distractors (0°: FR changes from 70% to 30%: −4.2±0.5 Hz, p<10^-9^; n=92 cells; Friedman test (p<10^-25^) with post-hoc Tukey HSD test; **Figure S3b**).

**Figure 3.**
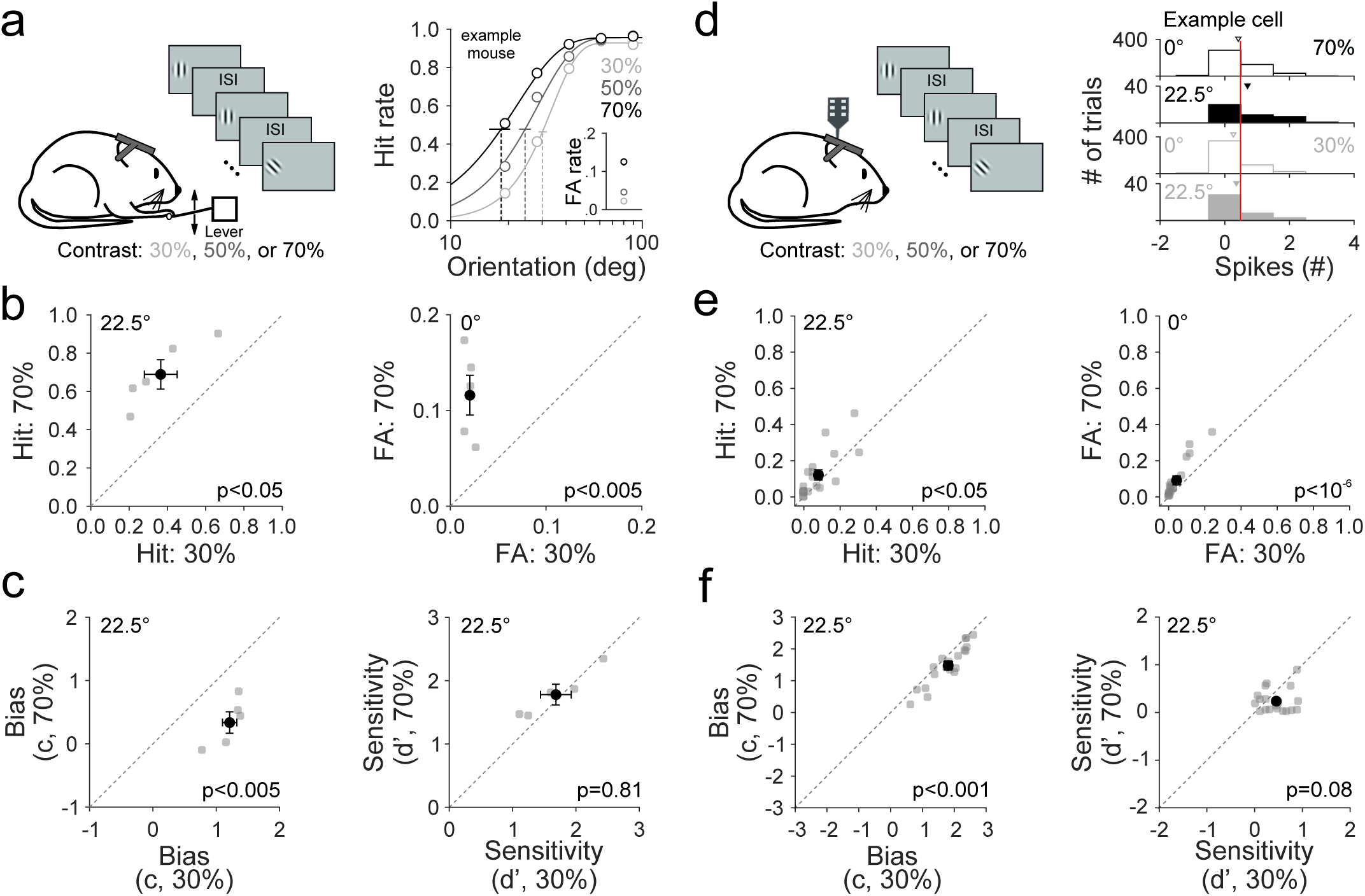
Decreasing stimulus contrast decreases both hit and FA rate and increases behavioral and neuronal bias. (**a**) Left: schematic of behavioral setup. Stimulus contrast is varied (30% (light gray), 50% (dark gray) or 70% (black)) on each stimulus presentation. Right: hit rate and FA rate (inset) for each contrast for an example mouse. Hit rates are fit with a Weibull function; vertical dotted lines are threshold, error is 95% confidence interval. (**b**) Comparison of hit (left, 22.5° target) and FA rate (right, 0° distractor) between two contrasts (70% vs 30%). Gray circles are individual mice and black circle is the mean of the population. Error bars are SEM across 5 mice. (**c**) Same as **b**, for bias (left) and sensitivity (right) at 22.5°. (**d**) Left: schematic of extracellular recording setup. Right: response distributions to distractor (0°, open bars) and target (22.5°, filled bars) stimuli of 30% (light gray) and 70% (black) contrast in an example cell. Triangles are the mean of the distribution. The criterion (vertical red line; see Methods) was determined for each cell and used to predict neuronal hit and false alarm rates for all contrasts. (**e-f**) Same as **b-c**, for predicted (**e**) hit and FA rate and (**f**) bias and sensitivity from the neuronal data (n = 19 cells, 4 mice).

Consistent with lower stimulus contrast driving lower firing rates, we find that decreasing stimulus contrast significantly reduces the animal’s hit rate (22.5° target: p<0.05; n=5 mice; one-way anova (p=0.05) with post-hoc Tukey HSD test; **Figure 3b**) and FA rate (70% vs. 30%: p<0.005; n=5 mice; one-way anova (p<0.005) with post-hoc Tukey HSD test). These changes in hit and FA rate drive a significant increase in bias (c for 22.5° target: 70% vs. 30%: p<0.005; n=5 mice; one-way anova (p<0.005) with post-hoc Tukey HSD test; **Figure 3c**) without a significant change in sensitivity (d’ for 22.5° target: p=0.81; n=5 mice; one-way anova).

As with optogenetic manipulation of neuronal activity, the effects of manipulating visual stimulus features on behavior are consistent with the changes in V1 activity. Lowering stimulus contrasts decreases both the predicted hit rate (22.5° target: 70% vs. 30%: p<0.05; Friedman test (p<0.01) with post-hoc Tukey HSD test; **Figure 3e**) and FA rate (0°: 70% vs. 30%: p<10^-5^; Friedman test (p<10^-6^) with post-hoc Tukey HSD test), resulting in an increase in measured bias (c for 22.5° target: 70% vs. 30%: p<10^-3^; Friedman test (p<10^-3^) with post-hoc Tukey HSD test; **Figure 3f**) without significantly changing the predicted sensitivity (d’ for 22.5° target: p=0.08; Friedman test). Thus, changes in the quality of sensory encoding through variation of visual stimulus properties can affect behavioral and neuronal measures of bias.

### Manipulation of adaptation state affects measures of bias

Varying stimulus contrast revealed that stimulus manipulations of sensory encoding can affect measured bias. In order to demonstrate the ubiquity of this phenomenon, we manipulated a different property of the task design that affects sensory encoding: inter-stimulus interval (ISI; 250, 500 and 750 ms; **Figure 4a**). Varying the ISI, like varying contrast, alters the strength of sensory responses, where shorter ISIs drive suppressive adaptation and lower firing rates (Clifford et al., 2007; Jin et al., 2018). Indeed, extracellular recordings revealed that adaptation significantly decreases the neuronal responses to distractors (0°: FR changes from 750 ms to 250 ms ISI: −3.9±0.8 Hz, p<10^-^ 8; n=74 cells; Friedman test (p<10^-9^) with post-hoc Tukey HSD test; **Figure S4b**), while slightly, but not significantly, decreasing responses to targets (22.5°: FR changes from 750 ms to 250 ms ISI: −2.4±0.8 Hz, p=0.17; n=74 cells; Friedman test; **Figure S4a**). While there is an asymmetric effect of ISI on targets and distractors (consistent with the stimulus specific effects of adaptation (Müller et al., 1999; Dragoi et al., 2000)), the net effect of adaptation is to reduce firing rates and this should decrease the optimal criterion and therefore increase bias.

**Figure 4.**
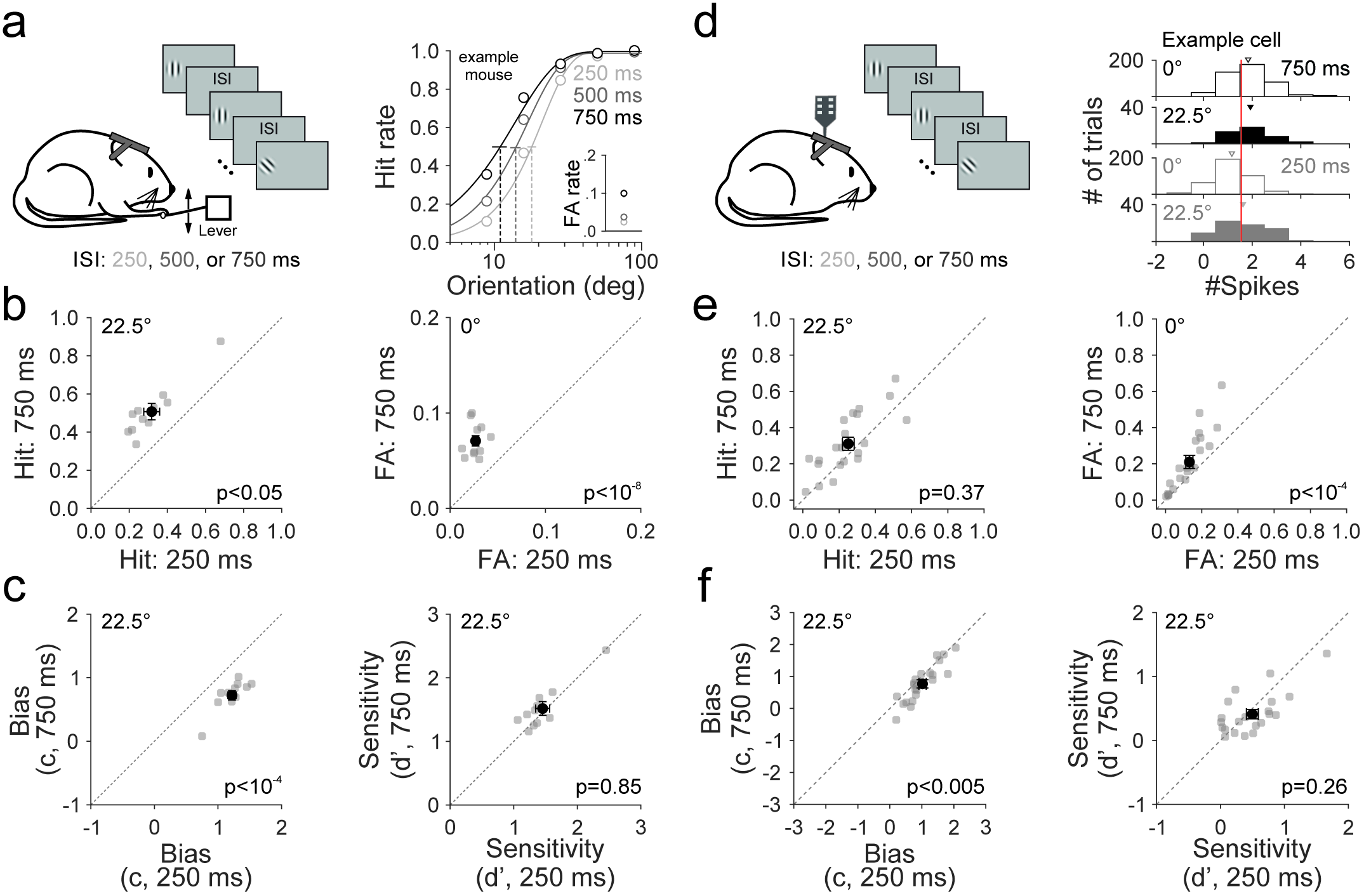
Adaptation decreases both hit and FA rate and increases behavioral and neuronal bias. (**a**) Left: schematic of behavioral setup. Inter-stimulus interval (ISI) is varied (250 ms (light gray), 500 ms (dark gray) or 750 ms (black)) on each stimulus presentation. Right: hit rate and FA rate (inset) for each ISI for an example mouse. Hit rates are fit with a Weibull function; vertical dotted lines are threshold, error is 95% confidence interval. (**b**) Comparison of hit (left, 22.5° target) and FA rate (right, 0° distractor) between two ISIs (750 vs. 250 ms). Gray circles are individual mice and black circle is the mean of the population. Error bars are SEM across 11 mice. (**c**) Same as **b**, for bias (left) and sensitivity (right) at 22.5°. (**d**) Left: schematic of extracellular recording setup. Right: response distributions to distractor (0°, open bars) and target (22.5°, filled bars) stimuli following 250 (gray) or 750 ms (black) ISI for an example cell. Triangles show the mean of the distribution. The criterion (vertical red line; see Methods) was determined for each cell and used to predict neuronal hit and false alarm rates for all ISIs. (**e-f**) Same as **b-c**, for predicted (**e**) hit and FA rate and (**f**) bias and sensitivity from the neuronal data (n = 21 cells, 4 mice).

Consistent with this prediction, decreasing the ISI decreases both hit rate (22.5° target: 750 ms vs. 250 ms: p<0.05 n=11 mice; one-way anova (p<0.05) with post-hoc Tukey HSD test; **Figure 4b**) and FA rate (750 ms vs. 250 ms: p<10^-8^; n=11 mice; one-way anova (p<10^-8^) with post-hoc Tukey HSD test). The decrease in both hit and FA rate support an increase in measured bias (c for 22.5° target: 750 ms vs. 250 ms: p<10^-4^; n=11 mice; one-way anova (p<10^-3^) with post-hoc Tukey HSD test; **Figure 4c**), without a coincident change in sensitivity (d’ for 22.5° target: p=0.85; one-way anova).

As with manipulating contrast, the behavioral effects of manipulating ISI are expected from the observed changes in neuronal activity recorded in V1. Using a fixed artificial decision criterion, the decreased responses to targets and distractors with decreasing ISI results in a significant decrease in the predicted FA rate (0°: 750 ms vs. 250 ms: p<10^-4^; Friedman test (p<10^-4^) with post-hoc Tukey HSD test; **Figure 4e**) and a slight, but not significant, decrease in the hit rate (22.5° target: p=0.37; Friedman test) resulting in an increase in measured bias (c for 22.5° target: 750 ms vs. 250 ms: p<0.005; Friedman test (p<0.005) with post-hoc Tukey HSD test; **Figure 4f**) without a change in sensitivity (d’ for 22.5° target: p=0.26; Friedman test). Thus, the effects of ISI on behavioral bias are consistent with the effects of ISI on sensory encoding. Thus, we have demonstrated that both direct optogenetic, and indirect stimulus-dependent, manipulations of sensory encoding affect both behavioral and neuronal measures of bias.

## Discussion

Signal detection theory is a standard approach for quantifying the sensory and cognitive contributions to perceptual decision-making. However, we provide both behavioral and neuronal evidence that measures of bias are sensitive to changes in sensory encoding. Directly manipulating neuronal excitability in V1 induced predictable changes in behavioral bias with comparatively little effect on sensitivity in the performance of an orientation discrimination task. Moreover, by varying either stimulus contrast or adaptation state, we also observed robust changes in bias. These results clearly demonstrate that changes in bias are not necessarily due to cognitive mechanisms, and conversely, that the lack of a change in sensitivity does not preclude effects on sensory encoding.

The optogenetic and stimulus manipulations applied in this study altered the quality of stimulus encoding. These manipulations each either increase or decrease neuronal responses to both targets and distractors, thereby increasing or decreasing the optimal criterion. Thus, the coincident change in both hit and FA rate are interpreted in SDT as a change in measured bias, even in the absence of a change in decision criterion (**Figure 1a**). Neuronal recordings in V1 confirmed that the optogenetic and stimulus manipulations shifted the target and distractor response distributions in the same direction, although not necessarily by the same amount. For instance, we find that V1 excitation increases the response to distractors slightly more than for targets (Modulation index: 0° vs. 22.5°: 0.48±0.05 vs. 0.37±0.06; p=0.07, n=83 cells; Wilcoxon signed rank test), likely due to the contribution of normalization circuits (Carandini and Heeger, 2012; Histed, 2018). These disproportionate changes in target and distractor distributions result in a change in both neuronal and behavioral sensitivity (**Figure 2**).

Importantly, measurements of bias from the neuronal activity clearly demonstrate that there can be changes in bias without changes in the decision criterion. We recorded from passively viewing mice to rule out the possibility that feedback from cognitive structures might influence the sensory responses. Moreover, in these analyses, we set and fix the decision criterion across conditions. While it is possible that the optogenetic and stimulus manipulations affect the animals’ decision criterion, we think it is unlikely. First, the optogenetic and stimulus conditions were varied on a presentation-by-presentation basis such that the animal could not predict the upcoming condition. Therefore, it is unlikely that the mouse could adjust its decision criterion on these short time scales. Even if optogenetic manipulations in V1 did change the decision criterion, the decision criterion would likely remain shifted for the immediately following stimulus after optogenetic termination within a trial. However we did not observe any changes in the behavior measures at the current stimulus when its preceding stimulus was suppressed or excited (c for 22.5°: V1 Stim_N-1_ suppression vs. Stim_N-1_ control: 0.9±0.2 vs. 0.8±0.2, p=0.51, n=4 mice; V1 Stim_N-1_ excitation vs. Stim_N-1_ control: 1.0±0.1 vs. 0.9±0.1, p=0.31, n=4 mice; paired t-test; **Figure S2**). Second, these manipulations do not significantly affect lapse rate across conditions (control vs. V1 suppression: p=0.13, n=4 mice, paired t-test; control vs. V1 excitation: p=0.18, n=4 mice, paired t-test; across contrasts: p=0.32, n=5 mice, one-way anova; across ISIs: p=0.97, n=11 mice, one-way anova).

However, if the animal were to compensate for the changes in sensory encoding by shifting its decision criterion, this could cancel the effects of sensory encoding on bias, making it seem as though there were no change in bias at all. Therefore, a lack of a change in bias does not guarantee a stable decision criterion. As we have shown, changes to sensory encoding that alter the target and distractor distributions in the same direction are commonplace. For instance, the classic gain-change effects of both spatial and feature attention on neuronal activity should drive changes in both sensitivity and bias (Treue and Maunsell, 1996; Treue and Martinez-Trujillo, 1999). In contrast, changes to sensory encoding that proportionally change target and distractor distributions in opposite directions, such that the optimal criterion is stable, are less common.

Notably, these manipulations are able to induce large shifts in bias in part because of the strategy that the mouse is using to perform the task (Jin et al., 2018). The circuits downstream of V1 are monitoring the total firing rates of a population of target-responsive sensory neurons. When the firing rate of this population exceeds some threshold, a change is detected. This can explain why increases in contrast, ISI or excitability of V1 neurons are often mistaken for target orientations and result in increased hit and FA rates. However, other decoding strategies that compute the estimated orientation from the population activity, for instance through a likelihood function, are also sensitive to manipulations of excitability in sensory cortex due to changes in certainty, and thus may also affect measured bias (Stocker and Simoncelli, 2006).

We find that the effects of optogenetic manipulation of neuronal activity in V1 on neuronal and behavioral bias go in the same direction. However, since we do not know the quantitative transform between sensory responses and behavior, these data cannot determine whether all of the changes in behavioral bias can be accounted for by changes in sensory encoding. Thus, while our optogenetic data is most consistent with a sensory role for V1, we cannot rule out some cognitive contributions. This reveals that combining optogenetics and SDT to dissociate the sensory and cognitive contributions to perceptual decision-making in distinct brain circuits is not straightforward. Realizing this confound, some groups have designed tasks to support the dissociation of sensory and cognitive contributions through SDT analyses. One such approach is to take advantage of the temporal separability between these processes. For instance, studies normally use pre-stimulus cues to bias the behavioral choice, but by adding a post-stimulus cue design one can better dissociate the effects of cue on sensory encoding and response bias (Bang and Rahnev, 2017). Other groups have taken advantage of clever stimulus design. For instance, using noisy stimulus sets to generate trial-by-trial variability enables experimenters to use regression-based approaches to measure stimulus sensitivity across conditions, and thereby dissociate of perceptual and response bias (Wyart et al., 2012; Kloosterman et al., 2018). Together, these approaches can be combined with optogenetics to determine the extent to which brain areas and circuits contribute to the various stages of perceptual decision-making.

## Methods

### Animals

All animal procedures conformed to standards set forth by the NIH, and were approved by the IACUC at Duke University. 23 mice (both sexes; 3-24 months old; singly and group housed (1-4 in a cage) under a regular 12-h light/dark cycle; C57/B6J (Jackson Labs #000664) was the primary background with up to 50% CBA/CaJ (Jackson Labs #000654)) were used in this study. *Pvalb-cre* (*tm1(cre)Arbr*, Jackson Labs #008069; n=15; PV::Cre), *VGAT-ChR2-EYFP* (*Slc32a1-COP4*H134R/EYFP*, Jackson Labs #014548; n=2) and Emx1-IRES-Cre (*tm1(cre)Krj*, Jackson Labs # 005628; n=6; EMX1::Cre) were crossed to C57/B6J mice for *in vivo* extracellular electrophysiology (n=11) and behavior (n=14) experiments. Note two of the mice (one PV::Cre and one Emx1::Cre) were used in both behavior and recording.

### Cranial window implant

Dexamethasone (3.2 mg/kg, s.c.) and Meloxicam (2.5 mg/kg, s.c.) were administered at least 2 h before surgery. Animals were anesthetized with ketamine (200 mg/kg, i.p.), xylazine (30 mg/kg, i.p.) and isoflurane (1.2-2% in 100% O_2_). Using aseptic technique, a headpost was secured using cyanoacrylate glue and C&B Metabond (Parkell), and a 5 mm craniotomy was made over the left hemisphere (center: 2.8 mm lateral, 0.5 mm anterior to lambda) allowing implantation of a glass window (an 8-mm coverslip bonded to two 5-mm coverslips (Warner no. 1) with refractive index-matched adhesive (Norland no. 71)) using Metabond.

The mice were allowed to recover for one week before habituation to head restraint. Habituation to head restraint increased in duration from 15 min to >2 h over 1-2 weeks. During habituation and electrophysiology sessions, mice were head restrained while either allowed to freely run on a circular disc (InnoWheel, VWR) or rest in a plastic tube.

### Visual stimulation

Visual stimuli were presented either on a 144-Hz (Asus) or 120-Hz (Samsung) LCD monitor, calibrated with an i1 Display Pro (X-rite), for electrophysiology and behavior experiments, respectively. The monitor was positioned 21 cm from the contralateral eye. Circular gabor patches containing static sine-wave gratings alternated with periods of uniform mean luminance (60 cd/m^2^). Visual stimuli for electrophysiology and behavior experiments were controlled with MWorks (http://mworks-project.org).

Three visual stimulus protocols were used for electrophysiology experiments in which we varied: 1) blue light (473 nm) stimulation of single stimulus presentations (each trial targeted with equal probability either: the distractor two stimuli before the target, the distractor before the target, the target, or no stimulation) and target orientations (22.5°, 45° and 90°; n=3 mice for each excitation and inhibition; **Figure 1-2**); 2) stimulus contrast (30, 50 and 70%) and target orientation (22.5° and 90°; n=5 mice **Figure 3**); and 3) number of distractor presentations (two to nine), inter-stimulus interval (ISI; 250, 500 and 750 ms) and target orientation (22.5°, 45° and 90°; n=4 mice; **Figure 4**). In the case that the stimulus properties were not varied, the default was six distractor presentations, a 250 ms ISI, 100% contrast. In order to maximize the contrast-dependence of neuronal responses, Protocol 2 used a 20° diameter gabor at a spatial frequency (SF) of 0.16 cyc/deg, to limit the contribution of increasing surround suppression with increasing contrast. Protocols 1 and 3 used a 30° gabor at a SF of 0.1 cyc/deg. All protocols had an inter-trial interval (ITI) of 4 s. All stimulus conditions that were varied on a trial-by-trial or presentation-by-presentation basis were randomly interleaved.

### Retinotopic mapping

Retinotopic maps generated from intrinsic autofluorescence or cortical reflectance (for *VGAT-ChR2-EYFP* mice). For intrinsic autofluorescence, the brain was illuminated with blue light (473 nm LED (Thorlabs) or a white light source (EXFO) with a 462 ± 15 nm band pass filter (Edmund Optics)), and emitted light was measured through a green and red filter (500 nm longpass); for cortical reflectance, the brain was illuminated with orange light (530 nm LED (Thorlabs)), and all of the reflected light was collected. Images were collected using a CCD camera (Rolera EMC-2, Qimaging) at 2 Hz through a 5x air immersion objective (0.14 numerical aperture (NA), Mitutoyo), using Micromanager acquisition software (NIH). Stimuli were presented at 4-6 positions (drifting, sinusoidal gratings at 2 Hz) for 10 s, with 10 s of mean luminance preceding each trial. Images were analyzed in ImageJ (NIH) to measure changes in fluorescence (dF/F; with F being the average of all frames) to identify primary visual cortex (V1) and the higher visual areas. Vascular landmarks were used to identify targeted sites (V1) for electrophysiology and optogenetics experiments.

### Viral injection

We targeted V1 in PV::Cre mice (n=4) for expression of Channelrhodopsin2 (ChR2) and in Emx1::Cre mice (n=6) for expression of Chronos. Dexamethasone (3.2 mg/kg, s.c.) was administered at least 2 h before surgery and animals were anesthetized with isoflurane (1.2-2% in 100% O_2_). The coverslip was sterilized with 70% ethanol and the cranial window removed. A glass micropipette was filled with virus (AAV5.EF1.dFloxed.hChR2.YFP (UPenn CS0384) or AAV9.hSyn.FLEX.rc.Chronos.GFP (Addgene 59056)), mounted on a Hamilton syringe, and lowered into the brain. 50 nL of virus were injected at 250 and 500 μm below the pia (30 nL/min); the pipette was left in the brain for an additional 10 minutes to allow the virus to infuse into the tissue. Following injection, a new coverslip was sealed in place, and an optical cannula (400 μm diameter; Doric Lenses) was attached to the cranial window above the injection site. Optogenetic behavioral experiments and electrophysiology experiments were conducted at least two weeks following injection to allow for sufficient expression.

### Extracellular electrophysiology

Electrophysiological signals were acquired with a 32-site polytrode acute probe (either A4x8-5mm-100-400-177-A32 (4 shanks, 8 site/shank at 100 μm spacing) or A1x32-Poly2-5mm-50s-177-A32, (1 shank, 32 sites, 25 μm spacing), NeuroNexus) through an A32-OM32 adaptor connected to a Cereplex digital headstage (Blackrock Microsystems). Unfiltered signals were digitized at 30 kHz at the headstage and recorded by a Cerebus multichannel data acquisition system (Blackrock Microsystems). Visual stimulation synchronization signals were also acquired through the same system via a photodiode directly monitoring LCD output.

On the day of recording, the cranial window was removed, and a small durotomy performed to allow insertion of the electrode into V1. A ground wire was connected via a gold pin cemented in a burrhole in the anterior portion of the brain. The probe was slowly lowered into the brain (over the course of 15 min with travel length of around 800 μm) until the most superficial recording site was in the brain and allowed to stabilize for 45 - 60 min before beginning recordings. For optogenetic stimulation in protocol 1, the optic fiber was held in place via an articulated arm (Flexbar, SKU: 14830) to allow light delivery (473 nm LED, Thorlabs) to the recording site. For V1 suppression, the mean light power was 0.28±0.02 mW (range: 0.1-0.4 mW); and for V1 excitation, the mean light power was 0.05±0.003 mW (range: 0.03-0.06 mW), matching the ranges that were used in the behavioral tests.

Of the 11 mice that were used for extracellular electrophysiology, 3 were previously trained in the orientation discrimination task, 3 were trained in a contrast discrimination task, and 5 were naïve.

### Behavioral task

Animals were water scheduled and trained to discriminate orientations in visual stimuli by manipulating a lever. The behavior training and testing occurred during the light cycle. We first trained mice to detect full-field, 90° orientation difference (target) from a static grating. Most mice (n=12) were trained with a 0° distractor; however, 2 mice were trained with a 45° distractor. On the initial days of training, mice were rewarded for holding the lever for at least 400 ms (required hold time) but no more than 20 s (maximum hold time). At the end of the required hold time, the grating changed orientation and remained horizontal until the mouse released the lever (or the maximum hold time expired). Typically, within two weeks of training, the mice began releasing the lever as soon as the target orientation appears. Once the animals began reliably responding to the target orientation, we added a random delay between lever press and target stimulus to discourage adoption of a timing strategy. Over the course of the next few weeks, the task was made harder by (in roughly chronological order): 1) increasing the random delay, 2) decreasing the target stimulus duration and reaction time window, 3) removing the stimulus during the ITI, 4) shrinking and moving the stimuli to more eccentric positions, 5) adding a mean-luminance ISI to mask the motion signal in the orientation change, and finally 6) introducing hard targets (range: 9-90°). Delays after errors were also added to discourage lapses and early releases.

In the final form of the task, each trial was initiated when the ITI (3s) had elapsed and the mouse had pressed the lever. Trial start triggered the presentation of a 100 ms static sinusoidal, gabor patch (30° in diameter, SF of 0.1 cycle/deg, positioned at an eccentricity of 30° - 40° in azimuth and 0° - 10° in elevation) followed by an ISI randomly selected on a presentation-by-presentation basis (250, 500 or 750 ms). For a subset of mice (n=5, **Figure 3**), the contrast of each presentation was also randomized (Michelson contrast: 30%, 50% and 70%); in these experiments the stimulus size was reduced to 20° and the SF increased to 0.16 cycle/deg, at 5°-15° in azimuth and 10° in elevation to 1) reduce the surround suppression and 2) compensate for the difficulty induced by low contrast and small size of the stimuli. The target orientation occurred with a random delay (flat distribution) after the first two presentations on each trial and the target orientation was randomly selected from a fixed set of values around each animal’s threshold. Mice received water reward if they released the lever within 100-650 ms (sometimes extended to 1000 ms) after a target occurred. However, for calculating hit and false alarm (FA) rate, we use a narrower reaction window (200-550 ms) to ensure that the majority of the releases in this window are due to stimulus driven responses and have independent reaction windows for adjacent stimuli with short ISIs.

For optogenetic stimulation (**Figure 1-2**), we delivered blue light to the brain though the cannula from a 473 nm LED (Thorlabs) or a 450 nm laser (Optoengine) and calibrated the total light intensity at the entrance to the cannula. The light power is titrated so that it does not induce significant changes in the lapse rate for both V1 suppression (lapse rate: control vs. V1 suppression: 0.08±0.01 vs. 0.10±0.02; p=0.13, n=4 mice, paired t-test) and V1 excitation (control vs. V1 excitation: 0.12±0.06 vs. 0.06±0.03; p=0.18, n=4 mice, paired t-test). For V1 suppression, the mean light power was 0.27±0.07 mW (range: 0.07-0.4 mW); and for V1 excitation, the mean light power was 0.06±0.02 mW (range: 0.02-0.1 mW, **Figure 2a-b**). On each trial, a single stimulus (either the distractor two stimuli before the target, the distractor before the target, the target, or the distractor after the target) was targeted with equal probability. The light was turned on around 30ms before the time of visual presentation onset for the duration of the stimulus (100 ms). Behavioral control was done with MWorks, and custom software in MATLAB (MathWorks).

Notably, there are overlapping animals in dataset of the optogenetic (**Figure 1-2**), contrast (**Figure 3**) and ISI manipulations (**Figure 4**). Below, we provided a table (**Table 1**) that describes the mice overlap and difference in time in collecting these datasets. Numbers (1-3) indicate the time sequence of the tasks that were tested and data was collected for each mouse, while 0 reflects no training on that task. Four mice were trained in a single task, 9 mice were trained on two tasks thus belonged to two datasets, and only 1 mouse was included in all datasets.

**Table 1.**
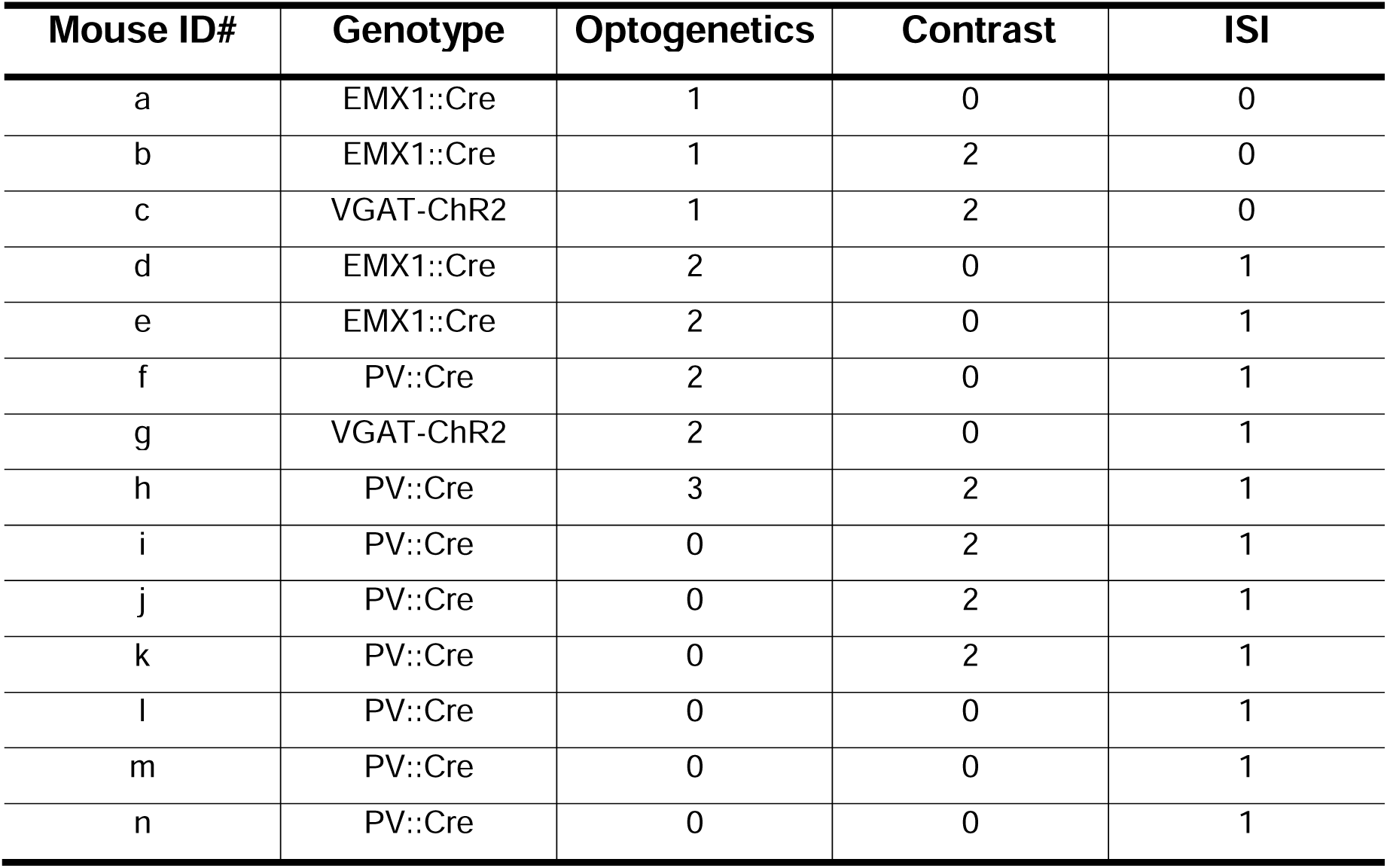
Mice overlap and timeline among three datasets

| Mouse ID# | Genotype | Optogenetics | Contrast | ISI |
| --- | --- | --- | --- | --- |
| a | EMX1::Cre | 1 | 0 | 0 |
| b | EMX1::Cre | 1 | 2 | 0 |
| c | VGAT-ChR2 | 1 | 2 | 0 |
| d | EMX1::Cre | 2 | 0 | 1 |
| e | EMX1::Cre | 2 | 0 | 1 |
| f | PV::Cre | 2 | 0 | 1 |
| g | VGAT-ChR2 | 2 | 0 | 1 |
| h | PV::Cre | 3 | 2 | 1 |
| i | PV::Cre | 0 | 2 | 1 |
| j | PV::Cre | 0 | 2 | 1 |
| k | PV::Cre | 0 | 2 | 1 |
| l | PV::Cre | 0 | 0 | 1 |
| m | PV::Cre | 0 | 0 | 1 |
| n | PV::Cre | 0 | 0 | 1 |

## Data processing

### Electrophysiology processing and analysis

Individual single units were isolated using the SpyKing CIRCUS package (http://spyking-circus.readthedocs.io/en/latest/). Raw data were first high pass filtered (> 500 Hz) and spikes were detected when a filtered voltage trace crossed threshold (9-13 median absolute deviations computed on each channel). A combination of density-based clustering and template matching algorithms were used to automatically cluster the spikes. The resulting clusters were then inspected and adjusted manually using a MATLAB GUI. Clusters with refractory period violations (< 2 ms, >1% violation) in the auto-correlogram and that were not stable across the whole recording session were discarded from the dataset. Clusters were combined if they met each of three criteria by inspection: 1) similar waveforms; 2) coordinated refractory periods in the cross-correlogram; 3) similar inter-spike interval distribution shape. Unit position with respect to the recording sites was calculated as the average of all site positions weighted by the waveform amplitude of each site. For V1-suppression or excitation experiments, we also quantified the similarity of the waveforms between control and optogenetic conditions using correlation coefficient (r) values. Because for majority of the cells, V1 suppression strongly reduce firing rate (**Figure S1c**) rendering few or even no spikes for analyzing waveforms, we extended the window starting from 200 ms before visual onset, end with 250 ms after visual offset. Signal and noise ratio of the trough value of the waveform shape was calculated as mean divided by SD across spikes. All of the subsequent analysis was performed in MATLAB.

Visually-evoked responses of each unit in V1 were measured based on average peri-stimulus time histograms (PSTHs, bin size: 20 ms) over repeated presentations (>25 trials) of the same stimulus. Response amplitudes were measured on a trial-by-trial basis: by subtracting the firing rate at the time of the visual stimulus onset from the value at the peak of the average PSTH within a window of 0-100 ms after the visual onset. However, in the case of V1 excitation, responses were measured by subtracting the baseline firing rate (value at visual onset, bin 0 ms, multiplied by 6) from the number of spikes during the visual presentation window (0-100ms, 6 bins). This is because the peak response latencies after V1 excitation were often shorter than the latencies of the visual responses in the control condition. “Responsive cells” were chosen as having statistically significant visually-evoked responses using a paired t-test to compare baseline responses (averaged over 0-100 ms before the visual onset) with visually-evoked responses (averaged over 0-100 ms after the visual onset; this analysis window excluded off-responsive units from analysis). For all protocols, we included cells that were significantly driven by either the first distractor stimulus or any of the target orientations. For V1 suppression experiments, we excluded cells that were significantly driven by the light stimulation. For protocol 2, this test was only performed for the highest contrast stimuli. Thus, we included 70/110 cells for V1 suppression; and 83/109 cells for V1 excitation; for protocol 2, we included 92/151 cells and for protocol 3, 74/100 cells were included; Modulation index (MI) of V1 excitation on neuronal responses (R) is calculated as:

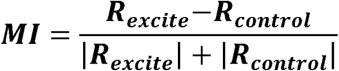

For calculation of predicted hit rate and FA rate: the distribution of single trial responses to the 22.5° target was compared to the distribution of responses to the distractor (0°, 3^rd^-6^th^ stimulus). For protocol 1 - V1 suppression, the artificial decision criterion for each cell was fixed as the mean of the responses to the suppressed distractor and the target in control. For protocol 1 - V1 excitation, the artificial decision criterion for each cell was fixed as the mean of the responses to the distractor in control conditions and the excited target. For protocol 2, the artificial decision criterion was fixed across all contrasts for each cell as the mean of the responses to the lowest contrast distractor (0°-30%) and the highest contrast target (22.5°-70%). For protocol 3, the artificial decision criterion was fixed across all ISIs for each cell as the mean of the responses to the most adapted distractor (0°-250ms ISI) and the most recovered target (22.5°-750 ms ISI). Thus, hit rate or FA rate across all conditions (either contrasts, ISIs or V1 suppression/excitation) were calculated as percentage of trials of the target or distractor responses that is higher than the artificial decision criterion, respectively.

Signal detection theory (Green and Swets, 1966) was applied to measure neuronal sensitivity (d’) and bias (c). For the extreme values of 0 and 1 for the predicted hit rate and FA rate, it is adjusted as follows to allow calculate sensitivity (d’) and bias (c): rates of 0 was replaced with 0.5/n, and rates of 1 was replaced with (n-0.5)/n, where n is the number of target or distractor trials (Macmillan and Kaplan, 1985; Stanislaw and Todorov, 1999). d’ and c were then calculated as follows:

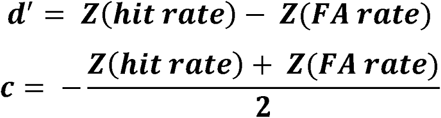

where Z is the inverse of the cumulative distribution function of the normal Gaussian distribution. To avoid confounds of directionality (since an increase in a positive d’ and a decrease in a negative d’ are both increases in sensitivity), only cells that had a positive d’ in the control condition (V1 suppression-47/70; V1 excitation-45/83), or across contrasts (19/92) and ISIs (21/74 cells) were included.

### Behavior processing and analysis

All behavioral processing and analysis were performed in MATLAB. All trials were categorized as either an early release, hit, or miss based on the time of release relative to target onset: responses occurring earlier than 100 ms after the target stimulus were considered early releases; responses occurring between 200 and 550 ms after the target were considered hits; failures to respond before 550 ms after the target were considered misses. Behavioral sessions were manually cropped to include only stable periods of performance and were further selected based on the following criteria: for protocol 1: 1) at least 40% of trials were hits; and 2) less than 50% of trials were early releases; for protocols 2&3: 1) at least 50% of trials were hits; and 2) less than 35% of trials were early releases. Based on these criteria, the data in **Figure 2 - V1 suppression** included 16 ± 3 (range: 8-19) sessions for each mouse with 4793 ± 706 trials (range: 3408-6695); **Figure 2 - V1 excitation** included 29 ± 13 (range: 3-58) sessions for each mouse with 8416 ± 3981 trials (range: 1551-18102); the data in **Figure 3** included 34 ± 13 (range: 11-75) sessions for each mouse with 8975 ± 3519 trials (range: 1017-23181), respectively; the data in **Figure 4** included 17± 3 sessions (range: 5-46) for each mouse with an average of 6348 ± 815 trials per mouse (range: 2593-11857).

Hit rate was computed from the number of hits and misses for each stimulus type:

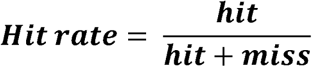

All distractor stimulus presentations were categorized as either a CR or a FA: responses occurring between 200 and 550 ms after a distractor stimulus were considered FAs; presentations where the mouse held the lever for at least 550 ms after the distractor stimulus were considered CRs. FA rate was computed from the total number of FAs and CRs in the session:

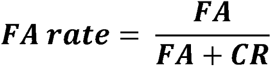

Hit and FA rate were used calculate behavioral d’ and c the same equations as were used to calculate d’ and c for the neuronal data. Since the detection threshold varies across mice and not all the mice have been sampled at exactly the same orientations such as 22.5°, the hit rate for 22.5° is extrapolated based on a Weibull function fitted from the psychometric curve for each mouse.

## Statistical analysis

Data were tested for normality using a Lilliefors test. While behavioral measures were normally distributed, electrophysiological measures of spike rates were not. Therefore, behavioral data were compared with either a t-test or ANOVA with post hoc Tukey HSD test for datasets with two and multiple groups, respectively. However, for the neuronal activity we used only non-parametric tests (Wilcoxon signed rank test and Friedman test with post hoc Tukey HSD test to compare two and multiple groups, respectively). Sample sizes were not predetermined by statistical methods, but are similar to other studies. Data collection and analysis were not performed blind to experimental conditions, but all visual presentation conditions in extracellular recording and behavior experiments are randomized.

## Data and code availability

All relevant data and code are available from the corresponding author upon reasonable request.

## Acknowledgements

We thank B. Gincley and J. Sims for assistance with behavioral training; K. Leonard, M. Fowler, J. Isaac and K. Murgas for surgical assistance; Z. Xu for assistance with software development; S. Lisberger, G. Field, C. Hull and members of the Hull and Glickfeld labs for helpful discussions and comments on the manuscript. This work was supported by an NIH Director’s New Innovator Award (DP2-EY025439), the Pew Biomedical Trusts, and the Alfred P. Sloan Foundation (L.L.G).

## Author Contributions

M.J. and L.L.G. designed the experiments. M.J. collected and analyzed the electrophysiology and behavior data. M.J. and L.L.G wrote the manuscript.

## Figure Legends

**Figure S1 – related to Figure 1.**
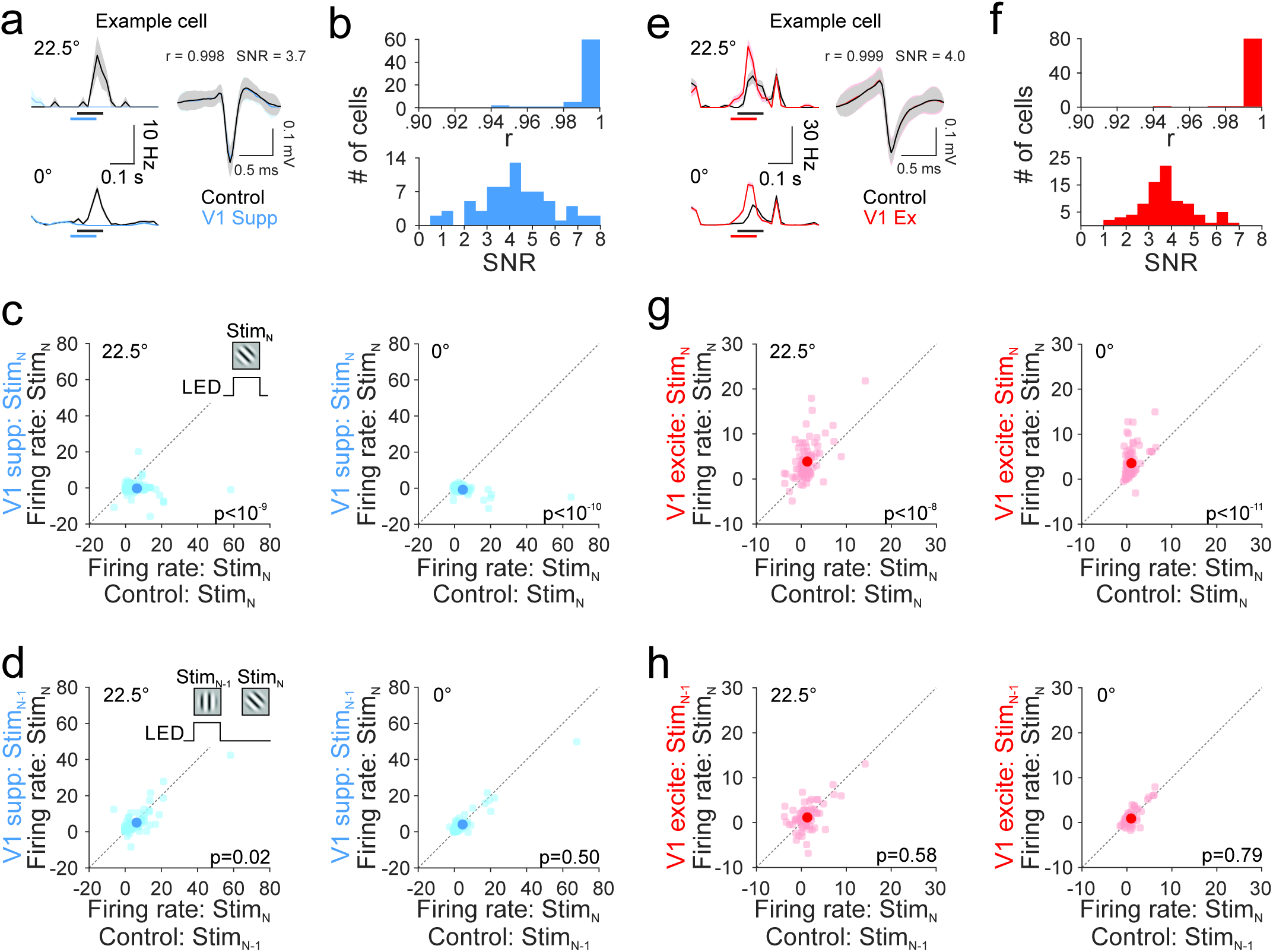
Optogenetically suppressing or exciting V1 decreases or increases neuronal responses to both targets and distractors. (**a**) Left: an example cell’s responses to 22.5° target (top) and 0° distractor (bottom) for control (black) and V1 suppression (blue). Shaded areas are SEM across trials. Right: mean waveform shapes for control and V1 suppression for the same cell in the left. Shaded areas are SD across spikes. Correlation coefficient (r) is shown to reveal the similarity in the waveform shapes between control and V1 suppression. Signal-to-noise ratio (SNR, mean/SD) of the trough value of the waveform is also shown. (**b**) Histogram of the correlation coefficient (r, top) and SNR values across all the cells (n=70 cells). (**c**) Comparison of neuronal responses (FR in Hz) to the 22.5° target (left) and 0° distractor (right) between control and V1 suppression (blue) on the current stimulus (Stim_N_). Light colors are individual cells and dark colors are the mean of the populations. Error bars are SEM across 70 cells (3 mice). (**d**) Comparison of neuronal responses to the 22.5° target (left) and 0° distractor (right) on Stim_N_ when the previous stimulus (Stim_N-1_) was suppressed vs. control. (**e-h**) Same as **a-d**, for V1 excitation (red, n=83 cells, 3 mice).

**Figure S2 – related to Figure 2.**
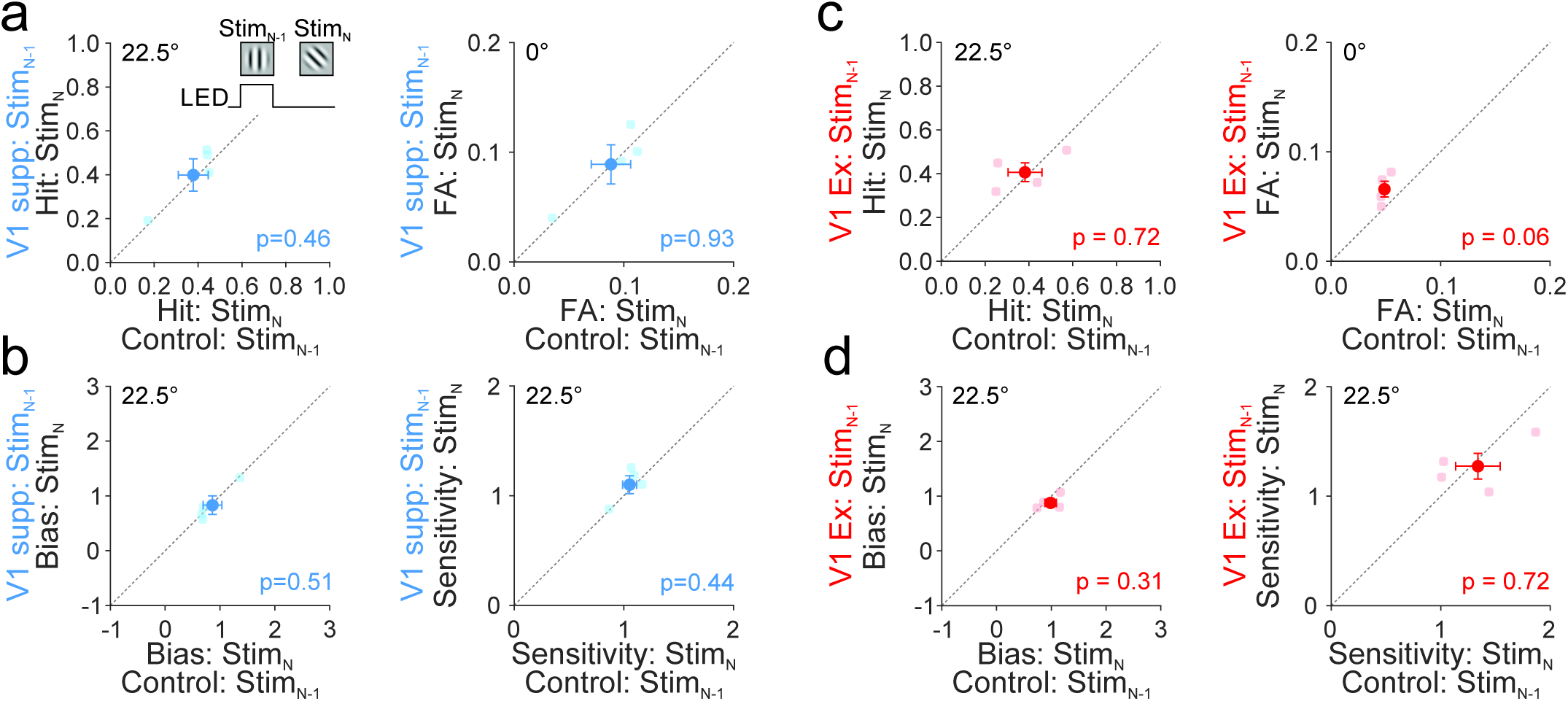
Lack of effects of Stim_N-1_ suppression or excitation on behavior measures at Stim_N_. (**a**) Comparison of hit rate (22.5°, left) and FA rate (0°, right) on Stim_N_ when the previous stimulus (Stim_N-1_) was suppressed (blue) vs. control. Light colors are individual mice and dark colors are the mean of the populations. Error bars are SEM across mice (n=4 mice). (**b**) Same as **a**, for bias (left) and sensitivity (right) measures for 22.5° target. (**c-d**) Same as **a-b**, for V1 excitation (red, n=4 mice).

**Figure S3 - related to Figure 3.**
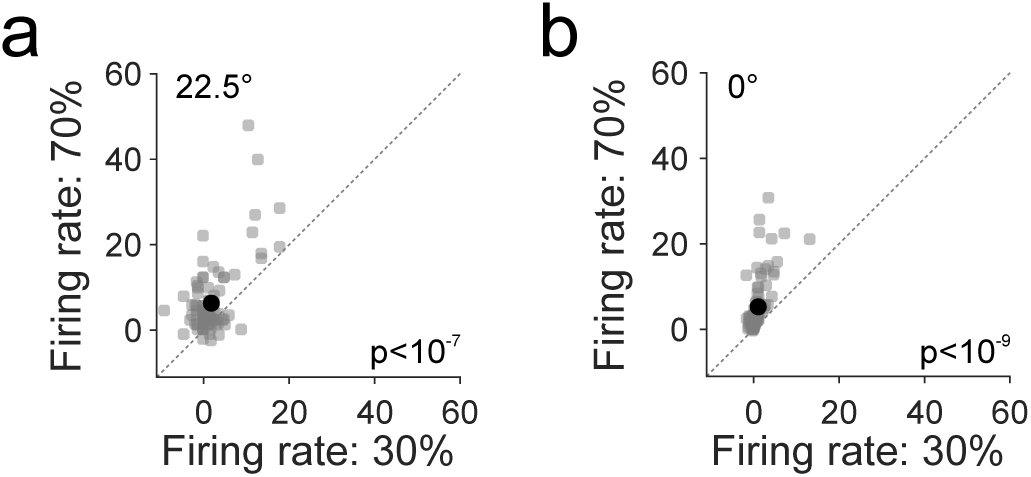
Reducing stimulus contrast reduces neuronal responses to both targets and distractors. (**a**) Comparison of neuronal responses (FR in Hz) to the 22.5° target between 30% and 70% contrasts. Gray circles are individual cells and black circle is the mean of the population. Error bars are SEM across 92 cells (4 mice). (**b**) Same as **a**, for responses to the 0° distractor.

**Figure S4 - related to Figure 4.**
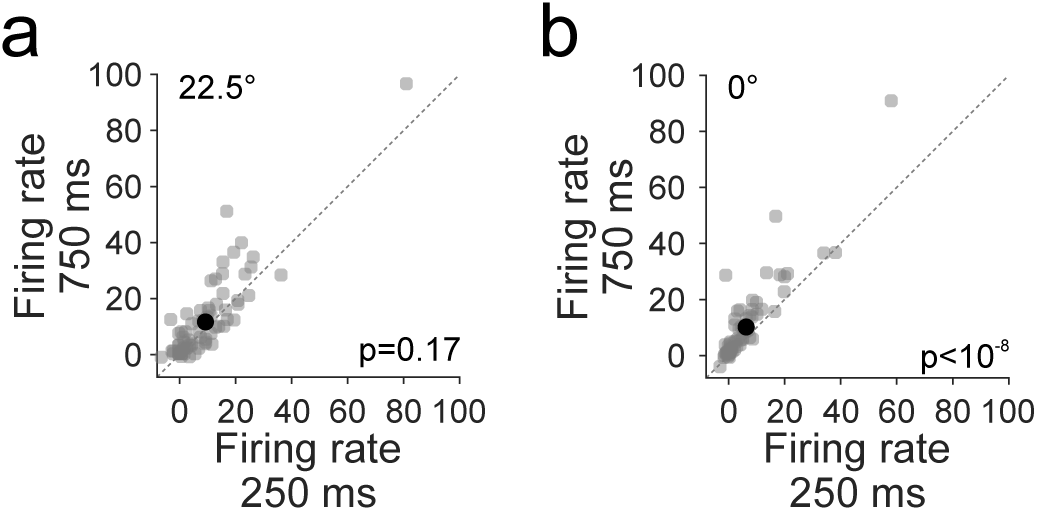
Adaptation reduces neuronal responses to both targets and distractors. (**a**) Comparison of neuronal responses (FR in Hz) to the 22.5° target after 750 or 250 ms ISIs. Gray circles are individual cells and black circle is the mean of the population. Error bars are SEM across 74 cells (4 mice). (**b**) Same as **a**, for responses to the 0° distractor.

